# Housing temperature influences exercise training adaptations in mice

**DOI:** 10.1101/651588

**Authors:** Steffen H. Raun, Carlos Henriquez Olguín, Iuliia Karavaeva, Mona Ali, Lisbeth L. V. Møller, Witold Kot, Josué L. Castro Mejía, Dennis Sandris Nielsen, Zach Gerhart-Hines, Erik A. Richter, Lykke Sylow

## Abstract

Exercise training is a powerful means to combat metabolic pathologies. Mice are extensively used to describe the benefits of exercise, but mild cold stress induced by housing temperatures may confound translation to humans. Thermoneutral housing is a strategy to make mice more metabolically similar to humans but its effects on exercise adaptations are unknown. Using voluntary wheel running, we show that thermoneutral housing blunted exercise-induced improvements in insulin action in muscle and adipose tissue. Moreover, thermoneutrality reduced the effects of training on energy expenditure, body composition, muscle and adipose tissue protein expressions, and the gut microbiome. The majority of these thermoneutral-dependent training adaptations could not be ascribed to a lower voluntary running volume. Thus, we conclude that organismal adaptations to exercise training in mice critically depend upon housing temperature. Our findings underscore the importance of housing temperature as an important parameter in the design and interpretation of murine exercise studies.

**Highlights:** - Housing at 30°C blunts several adaptations to exercise training in mice
- Exercise-sensitive protein induction is dampened at 30°C in skeletal muscle
- 30°C-housing blunts training-induced increase in insulin-stimulated glucose uptake
- Glucose tolerance is not improved by voluntary exercise training at 30°C housing
- Decreased running in 30°C housing is not due to overheating

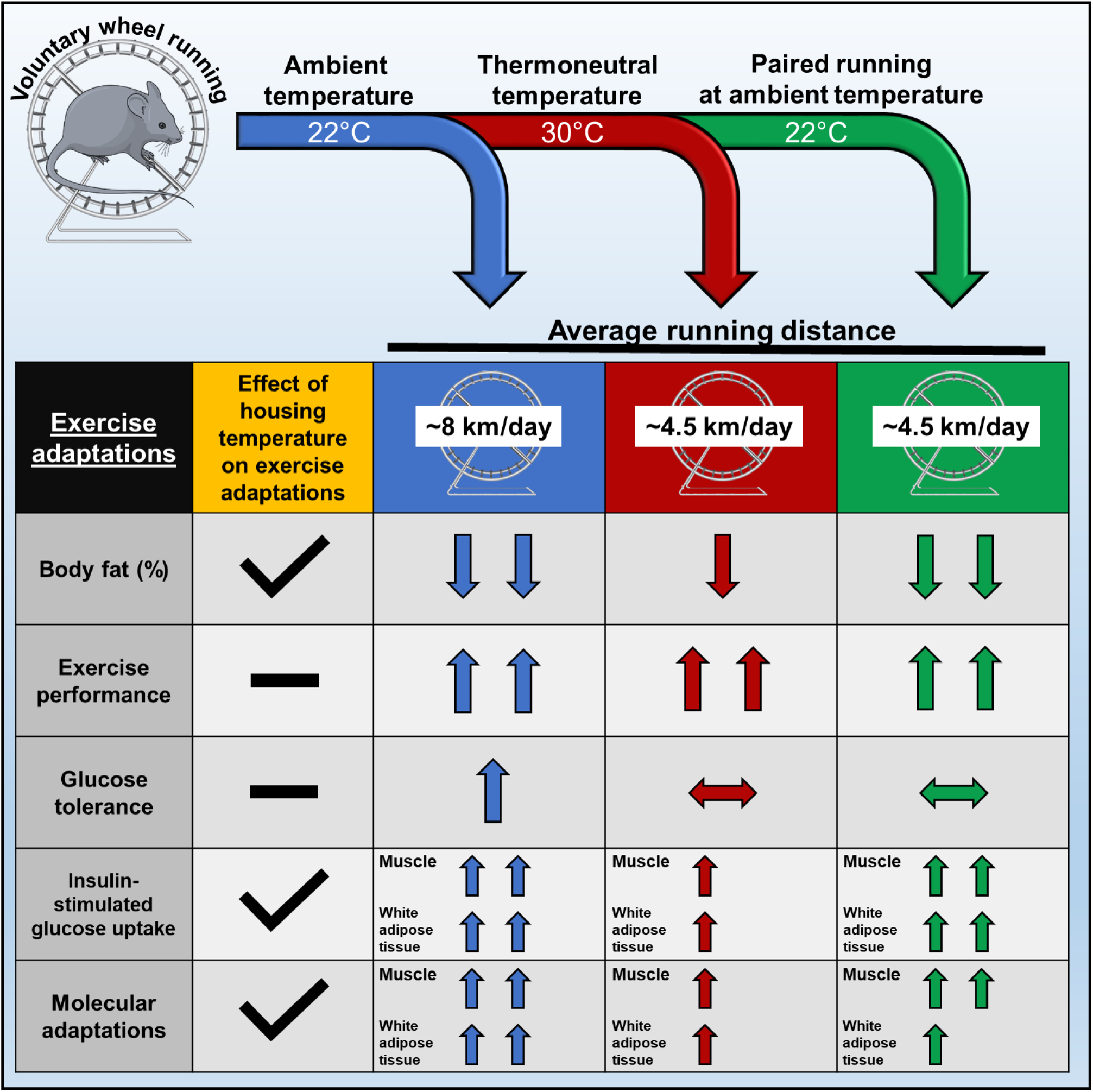
Graphical abstract.

## Introduction

Physical inactivity is a leading cause of morbidity and premature mortality worldwide^1,2^ and is associated with insulin resistance, obesity, and loss of muscle mass^3^. Regular exercise training is one of the most powerful means to combat such pathologies by eliciting health benefits on nearly all organ systems of the body^4–6^. Thus, much effort has been targeted towards understanding the underlying molecular mechanisms responsible for the adaptive responses to exercise training. In such studies, mice are extensively used as an experimental tool. However, failure to recognize that the laboratory mouse is housed under mild cold stress at ambient temperature^7,8^ may confound data interpretation and translatability to humans, who primarily live at thermoneutrality^9–11^. Thus, it might be time to rethink the optimal conditions for performing exercise studies in murine models, starting by housing mice at thermoneutral conditions to avoid chronic cold stress and its potential effects on adaptations to exercise training.

Mice prefer housing temperatures at 30°C compared to 20°C and 25°C^12^. Indeed, mice housed at ambient temperatures experience adverse effects on overall metabolic health^13^. They display sevenfold higher metabolic rate compared to humans^14^, have two-fold increased heart rate compared to mice housed at thermoneutrality^15^, show non-shivering thermogenesis due to increased sympathetic drive and activation of brown adipose tissue^8^, and exist right at the cusp of immune suppression ^16^. Experimentally these factors are obviously critical, although often underappreciated. Evidently, markedly dissimilar results have been obtained when testing the same processes in mice housed at different temperatures, including mitochondrial uncoupling^17^, whole-body glucose tolerance^18,19^ (although recently contradicted by^20^), inflammation^21^, immune responses^22^, atherosclerosis^23^, and cancer^24,25^. Instead, optimal housing conditions of mice to better mimic the metabolic rate of humans have been studied and discussed in recent years with temperatures from 27°C^26^ to 30°C^27,28^ being reported as the optimal housing temperature. Exercise markedly affects metabolism but to the best of our knowledge, the voluntary wheel running mouse model of exercise training has never been tested at thermoneutral conditions.

Here we undertake a detailed comparison of voluntary wheel running as an <u>E</u>xercise <u>T</u>raining model (ET) in mice housed at 22°C or 30°C. We show that housing temperature markedly influences the response to voluntary ET in skeletal muscle and adipose tissue, on glucose tolerance, insulin secretion, and the gut microbiome. Thus, our findings hold broad implications for assessing and interpreting systemic and molecular adaptive responses to voluntary training in mice at different housing temperatures.

## Results

### Exercise-induced changes in body composition and metabolic improvements are reduced at thermoneutrality

To elucidate the effect of housing temperature on exercise adaptations in mice, we housed mice at 22°C or 30°C with (Exercise Training; ET) or without (UnTrained; UT) free access to a running wheel for 6 weeks (excl. a 7-10 days temperature acclimatization period) (Fig 1A). This is one of the most commonly used training models for rodents often denoted *voluntary wheel running*. Body weight gain was similar between all groups during the 6 weeks intervention (Fig. 1B, Supplementary Fig. 1a) despite a 25% lower food intake in both ET and UT mice housed at 30°C, and a 30% increased food intake in both ET groups (Fig. 1C). At 30°C, ET attenuated gain in fat mass (−20%), while ET at 22°C, completely abolished fat mass gain (See Fig. 1D for body fat (%), See Suppl. Figure 1b for results in gram). For lean body mass, the opposite pattern was observed (see Fig 1E for LBM (%), see Suppl. Figure 1b for results in gram).

**Figure 1:**
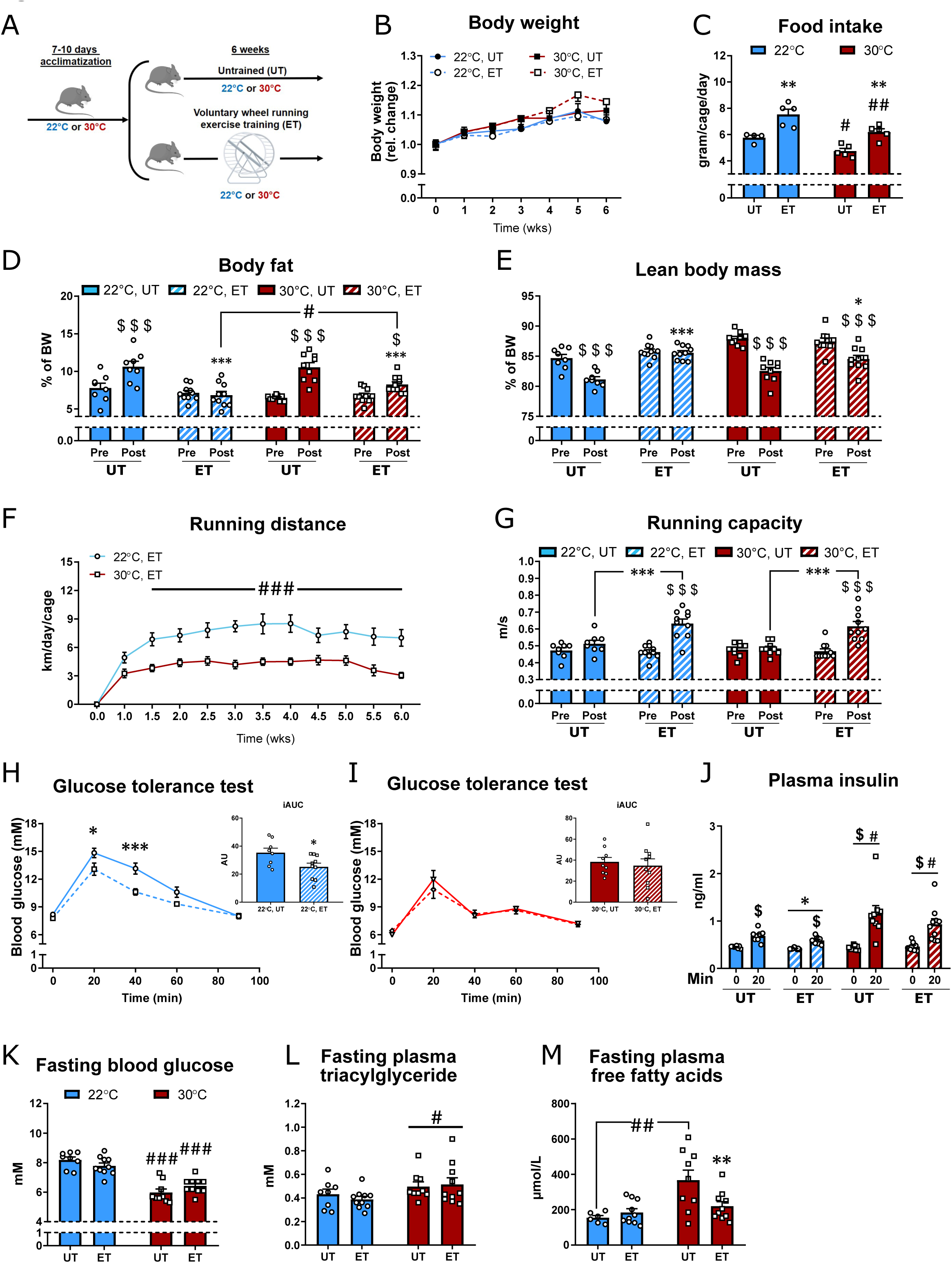
Voluntary exercise training in thermoneutrality induces smaller improvements on whole-body adaptations compared to 22°C. (A): graphic illustration of experimental training model. Mice were acclimatized to housing temperature before completing a 6 weeks voluntary wheel running exercise training (ET) intervention. (UT= untrained) (B): The effect of housing temperature and ET in 22°C and 30°C on bodyweight. n=8-10. (C): The effect of housing temperature and ET in 22°C and 30°C on food intake. Values are the average over 3 days of 3 different weeks from 4-5 cages. n=4-5. Effect of ET within temperature; ** p<0.01. Effect of temperature within UT or ET; # p<0.05, ## p<0.01. (D-E): The effect of housing temperature and ET in 22°C and 30°C on body fat (%) and lean body mass (%). n=8-10. Effect of time within group; $ p<0.05, $$$ p<0.001. Effect of temperature within ET-groups (post); # p<0.05. Effect of ET within temperature; * p<0.05, *** p<0.001. (F): Running distance per day in 22°C and 30°C, respectively. n=10. Effect of temperature; ### p<0.001. (G): Exercise capacity before (Pre) and after (Post) the training intervention. n=8-10. Effect of time within group; $$$ p<0.001. Effect of ET within temperature; *** p<0.001. (H-I): Effect of ET at 22°C and 30°C on glucose tolerance. n=8-10. Effect of ET on blood glucose response; * p<0.05, *** p<0.001. Effect of ET on iAUC; * p<0.05. (J): Effect of ET at 22°C and 30°C on glucose-stimulated insulin secretion at time 0min and 20min. n=8-10. Effect of time within group; $ p<0.05. Effect of temperature within UT or ET-groups (post); # p<0.05. Effect of ET within temperature; * p<0.05. (K): Fasting blood glucose after 5 hrs. of fasting. n=8-10. Effect of temperature within UT or ET; ### p<0.001. (L-M) The effect of housing temperature and ET in 22°C and 30°C plasma triglyceride and free fatty acids. n=8-10. Effect of temperature; # p<0.05, ## p<0.01. Effect of ET within temperature; * p<0.05. Data are presented as mean ± SEM incl. individual values where applicable.

Throughout the intervention, mice housed at 30°C ran 60% of the distance completed by mice housed at 22°C (Fig. 1F). Thermoneutral housing (30°C) was recently shown to decrease exercise performance after just 7 days^29^. In the current study, no effect of 6 weeks of thermoneutral housing could be detected on exercise performance in UT mice and in fact, ET elicited the same improvements (+35%) in maximal running speed (Fig. 1G), despite the 40% lower training volume at 30°C.

Having established that housing temperature altered ET adaptations on body composition but not exercise performance, we next asked whether housing temperature would affect the metabolic benefits to voluntary ET. Glucose tolerance improves following voluntary wheel running in mice^30–32^, but this has to the best of our knowledge only been tested for mice housed at 22°C. In contrast to ET at 22°C (Fig. 1H), ET at thermoneutral conditions did not improve glucose tolerance (Fig. 1I). This was despite a reduction in fat mass at both temperatures and improvements in running capacity by ET similar to what was seen at 22°C (Fig. 1D and G). We note that blood glucose during the glucose tolerance test (GTT) was noticeably lower in UT 30°C mice where it peaked at 12mM compared to UT 22°C mice where blood glucose peaked at 15mM. That suggests that the 30°C housing condition *per se* improves glucose tolerance in mice that could not be further improved by ET. In fact, glucose tolerance was similar between UT 30°C and ET 22°C housed mice (comparing Fig. 1H and I), which is in agreement with the notion that the standard control mouse at 22°C is less active and may be metabolically challenged^13^. In contrast to blood glucose, the 4-hour fasted plasma insulin concentration was similar between all experimental groups (Fig. 1J). Plasma insulin was reduced by ET in 22°C housed mice indicating improved insulin sensitivity, which was not observed in thermoneutrally housed ET mice (Fig. 1J). Interestingly, glucose-stimulated plasma insulin was 40% higher in mice housed at 30°C compared to 22°C, suggesting that housing temperature significantly affects β-cell function and/or sensitivity and/or insulin clearance (Fig. 1J). This was supported by higher HOMA-β in both UT and ET mice housed at 30°C (Suppl. Fig. 1c). HOMA-IR indicated improved insulin sensitivity by ET at 22°C (p=0.05), while mice in 30°C generally exhibited lower HOMA-IR (Suppl. Fig. 1d). Thermoneutrally housed mice had fasting blood glucose of ~6mM, while 22°C housed mice had ~8mM and these were unaltered by ET (Fig. 1K). The plasma triglyceride concentration (fasted) was 25% higher in 30°C housed mice compared to mice housed at 22°C with no effect of ET (Fig. 1L). In UT mice, plasma free fatty acids (fasted) were 135% higher at 30°C than in 22°C, where ET led to a reduction only in 30°C (Fig. 1M).

### Thermoneutral housing lowers energy expenditure and metabolic fluctuations in exercise-trained mice

To further elucidate the impact of temperature on whole body adaptations to exercise training, we subsequently conducted a series of experiments in metabolic chambers. We first sought to investigate to what extent and how rapid the metabolism of mice changes when increasing housing temperature from 22°C to 30°C. Slowly raising the temperature over ~3 hours caused a rapid drop in oxygen uptake (VO_2_; Suppl. Fig. 1e) and respiratory exchange ratio (RER; Suppl. Fig. 1h) within 6 hours of temperature change. The change in temperature led to a decrease (−45%) in energy intake (Suppl. Fig. 1f) without any changes in habitual activity (Suppl. Fig. 1g). These findings illustrate that housing temperature robustly and rapidly alters mouse metabolism. These results align with previous reports ^9,10,27,33^ and underline the metabolic challenges that are imposed on mice housed at room temperatures.

We next sought to determine the effects of temperature on whole body metabolism in mice already trained for 5 weeks (at 22°C or 30°C) by placing them in the metabolic chambers with or without access to a running wheel. Voluntary ET increased nightly VO_2_ compared to <u>UT</u> mice in both temperatures, but the effect of nighttime in ET was 60% higher at 22°C compared to 30°C housed mice (Fig. 2A and B). As expected, RER showed diurnal rhythm at 22°C, where RER during the day was reduced in ET compared to UT mice (Fig. 2C). When housed at 30°C, RER was similar between day and night in untrained mice, in contrast with previous reports ^20,27^, with no effect of ET on resting RER (Fig. 2C). Chronically housing mice at thermoneutrality increased habitual (+90%) activity during the night in UT mice (Fig. 2D) underlining the importance of adequate acclimatization time during temperature changes, as acute temperature did not change habitual activity (Suppl. Fig. 1g). Increased habitual activity was despite lower food intake compared to UT mice at ambient temperature (Suppl. Fig. 1i). This contrasts with the nightly running volume that tended to remain lower (p=0.064, −35%, Fig. 2E) in mice housed at 30°C compared to ambient temperature, while running volume during daytime was increased at 30°C (Fig. 2F). Overall mice housed at 30°C ran 35% less than when housed at 22°C (p=0.06, Fig. 2G) accompanied by a tendency to a lower maximal (p=0.054, Fig. 2H) and decreased average (Fig. 2I) running speed. To test if reduced running volume could be due to over-heating during exercise, we measured core temperature during the day (when the mice are resting/inactive) and in the early dark period (when the mice are running the most). UT mice displayed 0.6°C lower core temperature during the inactive period when housed at 22°C compared with 30°C, highlighting the mild cold stress inflicted by 22°C housing (Fig. 2J). Core temperature increased to the same absolute values in all groups during the dark period (Fig. 2J). Thus, reduced running of mice housed at 30°C is likely not due to overheating.

**Figure 2:**
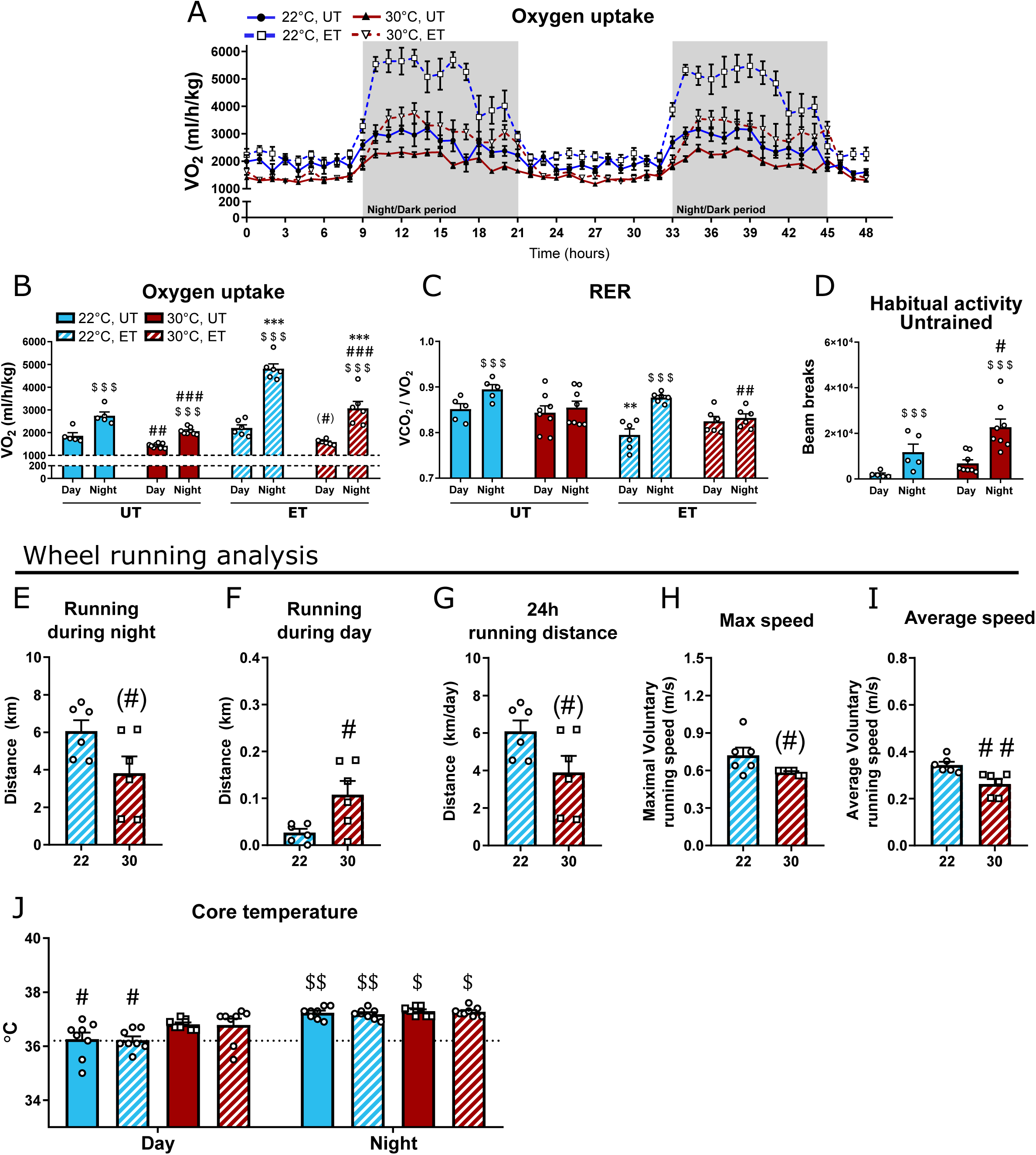
Housing temperature markedly affects the metabolic responses to exercise training in mice. (A-C): VO_2_ and RER in untrained (UT) and voluntary wheel running exercise trained (ET) mice at 22°C and 30°C housing. n=5-8. Effect of time within group; $$$ p<0.001. Effect of temperature within time of day; # p<0.05, ## p<0.01, ### p<0.001. Effect of ET within temperature; ** p<0.01, *** p<0.001. (D): Habitual activity (2 consecutive days) in UT mice after 6 weeks temperature acclimatization. n=5-8. Effect of time within group; $$$ p<0.001. Effect of temperature within time of day; # p<0.05. (E-I): Effect of temperature on wheel running distance, maximum and average speed. n=6-7. Effect of temperature; # p<0.05, ## p<0.01, (#) p<0.1. (J): Core temperature was measures at day (light period) and night (dark period) time via rectal thermometer, Effect of time, day vs. night; $ p<0.05, $$ p<0.01. Effect of temperature within day; # p<0.05. Data are presented as mean ± SEM incl. individual values where applicable.

Overall, mice in 30°C displayed lower energy consumption, lesser improvements in body composition, as well as no improvement in glucose tolerance following the standard laboratory exercise model of voluntary wheel running, despite similar improvements in running performance.

### Adaptations in glucose tolerance, but not performance or body composition, are ascribed to training volume

Having established remarkable differences in exercise training adaptations with this model in mice housed at different temperatures, we next sought to determine if these alterations were due to the lower running volume in 30°C housed mice. Thus, we restricted voluntary running in mice housed at 22°C (Fig. 3A) to mimic the training volume of thermoneutrally housed mice (“paired 22°C ET”, Fig. 3B).

**Figure 3:**
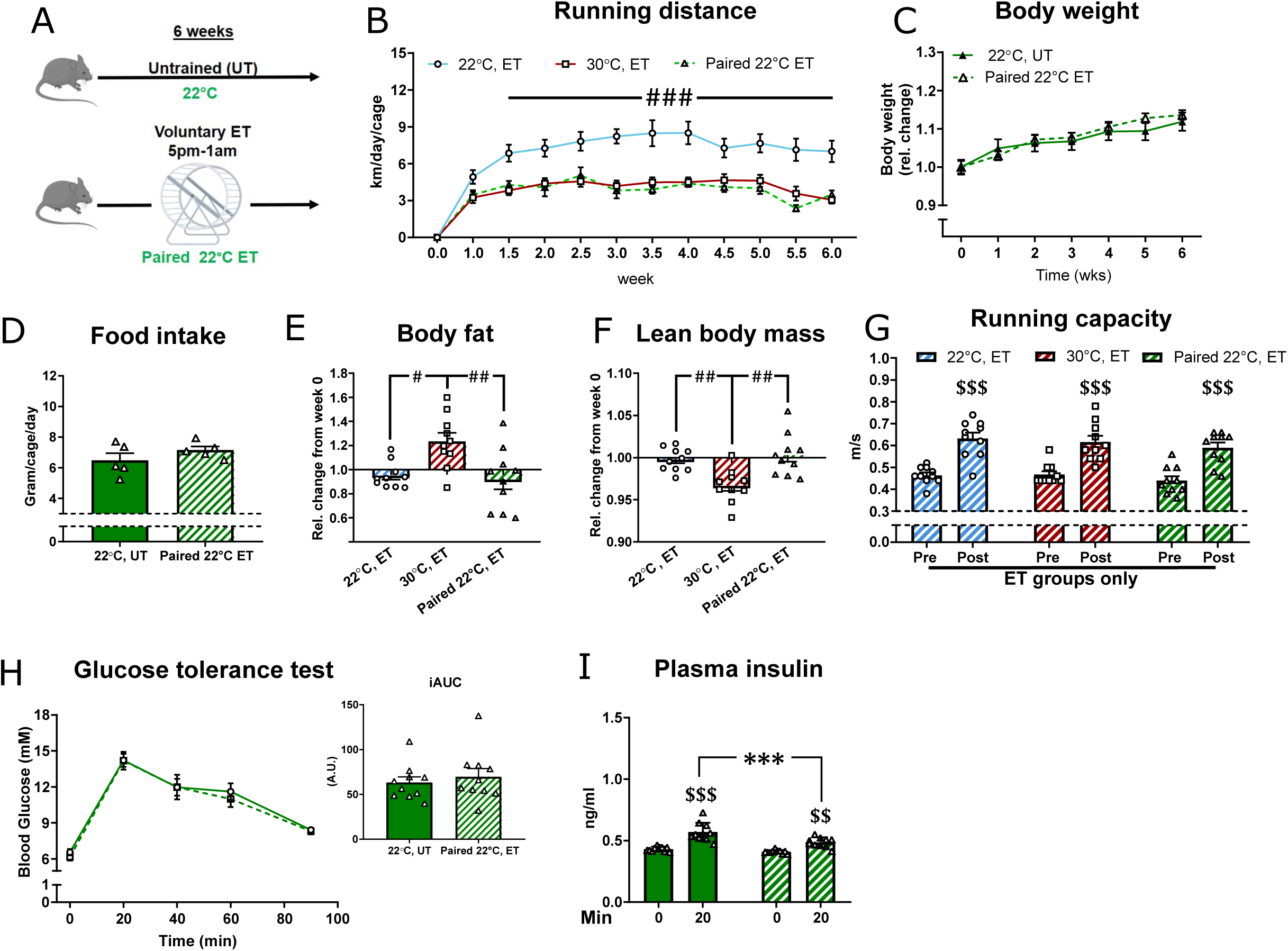
Training volume cannot account for the differences in exercise training responses in different housing temperatures. (A): Graphical illustration of the paired 22°C voluntary wheel running exercise training (ET) intervention. (B): Running distance per day in 22°C, 30°C, and paired 22°C, respectively. n=5-10. Effect of temperature; ### p<0.001. (C): The effect of paired 22°C ET on bodyweight. n=10. (D): The effect of paired 22°C ET on food intake. Values are the average over 3 days of 3 different weeks from 5 cages. (E-F): The relative change in body fat (%) and lean body mass (%) from before to after the ET intervention in 22°C, 30°C, and paired 22°C. n=10. Effect between the groups as indication with lines; # p<0.05, ## p<0.01. (G): Running capacity before and after the ET intervention in 22°C, 30°C, and paired 22°C. n=10. Effect of time within group; $$$ p<0.001. (H): Effect of paired 22°C ET on glucose tolerance. n=10. (I): Effect of paired 22°C ET on glucose-stimulated insulin secretion at time 0min and 20min. n=10. Effect of time within group; $$ p<0.01, $$$ p<0.001. Effect of ET; *** p<0.001. Data are presented as mean ± SEM incl. individual values where applicable.

Body weight (Fig. 3C) and food intake (Fig. 3D) were unaffected by paired 22°C ET. Similar to 22°C housing (Fig. 1D), paired 22°C ET reduced fat mass gain (Fig. 3E). As such, mice training at 30°C showed higher body fat (Fig. 3E) and lower lean body mass relative to body weight (Fig. 3F) compared to mice training at 22°C. Thus, housing temperature altered ET adaptations on body composition independent of training volume. Alongside these observations, exercise capacity was improved to the same extent by paired 22°C ET when compared to both ET at 30°C and 22°C (Fig. 3G). In contrast, paired 22°C ET did not improve glucose tolerance (Fig. 3H), suggesting that the differences between ET effects on glucose tolerance at different housing temperatures could be ascribed to training volume. However, plasma insulin concentrations were 15% lower in the paired ET compared to the UT group, suggesting that paired ET did in fact increase insulin sensitivity regardless of the lower running volume (Fig. 3I). No change in HOMA-β or HOMA-IR was observed (Suppl. Fig. 2).

These data indicate that, apart from glucose tolerance, which could to some extent be ascribed to running volume, there is a direct effect of housing temperature on ET-induced body composition adaptations.

### Thermoneutral housing prevents the improved insulin action in skeletal muscle following ET, independently of running volume

We next investigated if housing temperature would affect the ET-induced adaptations on insulin action and glucose uptake in skeletal and cardiac muscle.

Insulin was injected in the retro-orbital vein in anaesthetized mice. Insulin caused a drop of blood glucose that was similar between all experimental groups (Fig. 4A, B, and C). Plasma insulin concentration was 1.8 ng/ml at tissue harvest, 10 min following the insulin injection, in all groups (Suppl. Fig. 3a).

**Figure 4:**
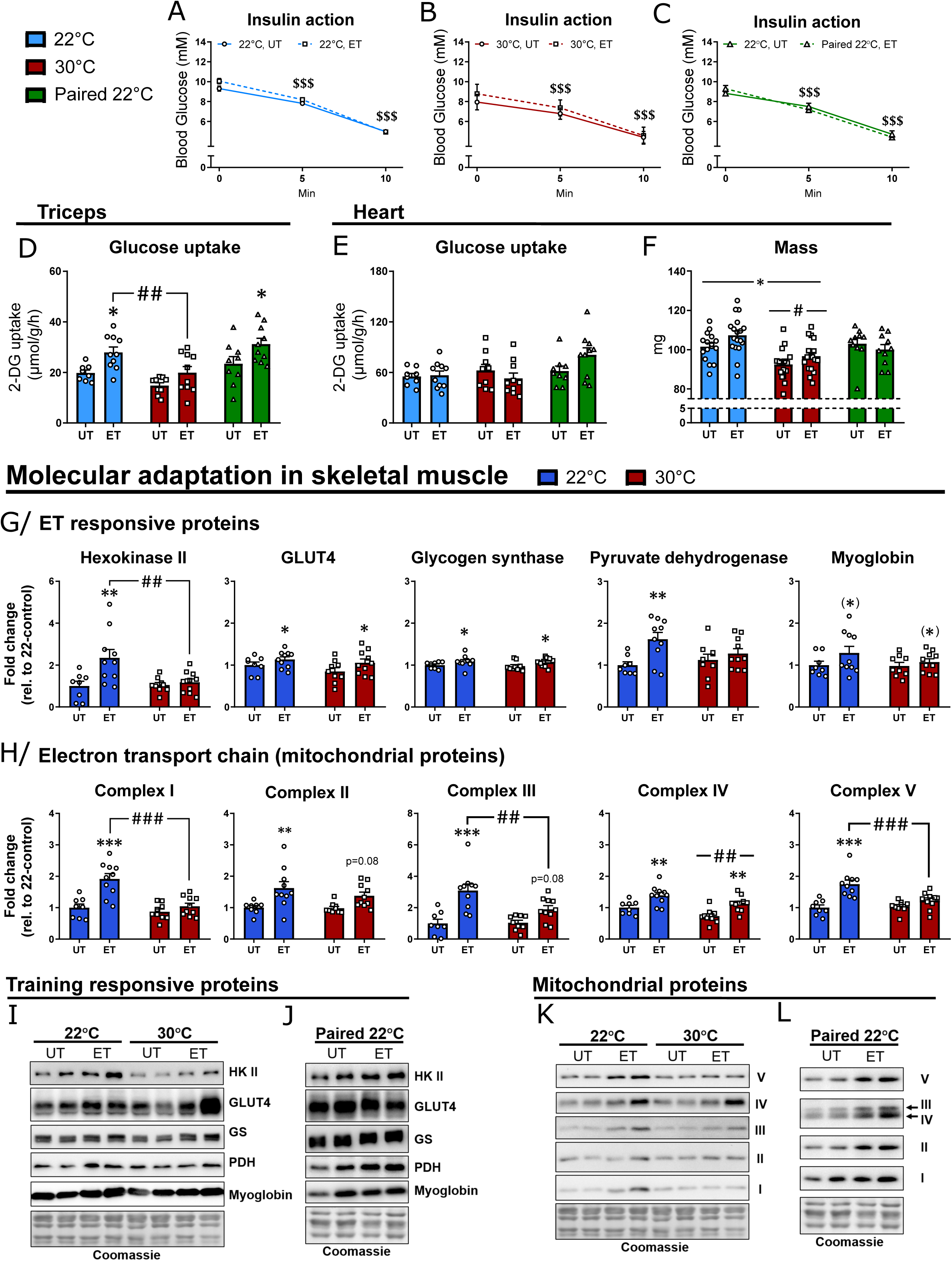
Skeletal muscle shows lesser improvements in insulin action and molecular adaptations after exercise training at thermoneutrality. (A-C): Effect of retro-orbital insulin injection (0.3U/kg) on blood glucose in 22°C (A), 30°C (B), and paired 22°C (C), untrained (UT) and voluntary wheel running exercise trained (ET). n=10-14. Effect of time (insulin); $$$ p<0.001. (D-E): Effect of ET on insulin-stimulated glucose uptake in 22°C, 30°C, and paired 22°C in skeletal muscle (m. triceps brachii) (D) and cardiac muscle(E). n=8-10. Effect of ET within temperature; * p<0.05. Effect of temperature within ET groups; ## p<0.01. (F): Effect of ET on cardiac muscle weight in 22°C, 30°C, and paired 22°C (UT and ET groups). n=10-18. Main-effect of ET; * p<0.05. Main-effect of temperature; # p<0.05. (G-H): Effect of ET on training responsive proteins (G) and subunits of the electron transport chain of the mitochondrion (H) in triceps muscle at 22°C and 30°C. n=8-10. Effect of ET within temperature; * p<0.05, ** p<0.01, *** p<0.001. Effect of temperature within UT or ET groups; ## p<0.01, ### p<0.001. (I-L): Representative blots of proteins investigated in (G-H) and paired 22°C ET (J and L). Quantitative bar-blots of J and L can be seen in Appendix. Fig. 3. Data are presented as mean ± SEM incl. individual values where applicable.

Despite no apparent effect of ET on the change in whole body blood glucose levels during insulin stimulation, insulin-stimulated glucose uptake was increased by ET in skeletal muscle (m. triceps brachii (triceps), +40%, Fig. 4D) at 22°C. This effect was not observed following ET at 30°C (Fig. 4D). This lack of ET-induced increase in muscle insulin-stimulated glucose uptake was ascribed to housing temperature, as the paired 22°C ET group exhibited increased insulin-stimulated muscle glucose uptake (+35%, Fig. 4D), in spite of the reduced running volume. Similar results were observed in quadriceps muscle (Suppl. Fig. 3b). Neither housing temperature nor ET affected insulin-stimulated glucose uptake in the heart (Fig. 4E). This was despite similar cardiac hypertrophy (+5%) after ET at both housing temperatures and a 9.5% reduction in heart mass in 30°C housed mice (Fig. 4F). However, the mass of the heart was unaffected by ET in the paired 22°C ET group. Basal glucose uptake (Suppl. Fig. 3c) and importantly, 2-deoxy-glucose (^3^H) tracer activity were similar between all groups (Suppl. Fig. 3d). These results show that thermoneutral housing prevents the improved insulin action in skeletal muscle following ET, independently of running volume.

### Thermoneutrality alters the molecular adaptations to ET in skeletal muscle without affecting canonical insulin signaling

Major molecular adaptations occur in skeletal muscle in response to exercise training, but such responses have to the best of our knowledge in mice only been shown at ambient temperature. We therefore determined the molecular responses to voluntary ET on known training responsive proteins. In triceps muscle, GLUT4 (+20%), glycogen synthase (GS; +10%) and myoglobin (+20%) all increased with voluntary ET irrespective of housing temperature (Fig. 4G). In contrast, hexokinase (HK) II (+135%) and pyruvate dehydrogenase (PDH) (+60%) increased following ET only in 22°C housed mice (Fig. 4G, see 4I for representative blots). In the paired 22°C ET mice, ET increased HKII (30%), GS (10%, p=0.07), PDH (+45%), while no changes in GLUT4 and myoglobin were observed (see representative blots in Fig. 4J, for bar plots see Suppl. Fig. 3g).

It is well known that mitochondrial content increases with exercise training. To our surprise, we observed reduced response in four of the five complexes of the electron transport chain (ETC) following ET of 30°C housed mice compared to 22°C (Fig. 4H, see 4K for representative blots). The fact that housing temperature affects mitochondrial adaptations was confirmed by the paired 22°C ET mice, where all complexes increased after the intervention (see representative blots in Fig. 4L, for bar plots see Suppl. Fig. 3g).

We observed no effect of ET or temperature on protein expression of any of the above described proteins in heart muscle (Suppl. Fig. 3e). In addition, during insulin stimulation, no major changes in canonical insulin signaling were observed with ET or housing temperature in any of the analyzed muscles (Suppl. Fig. 3f). Housing temperature-induced changes in ET response observed in insulin-stimulated glucose uptake in skeletal muscle could therefore not be ascribed to altered intracellular insulin signaling, but rather changes in expression of glucose-handling proteins.

Collectively these data demonstrate that the ability of ET to increase insulin-stimulated glucose uptake and protein expression of key training responsive proteins in skeletal muscle were lost or markedly diminished when the mice were housed at 30°C, and this was not due to lower training volume.

### Temperature-dependent adaptation of adipose tissue is not influenced by exercise training

As white (WAT) and brown (BAT) adipose tissue are responsive to temperature^34,35^ as well as ET at ambient temperature^36,37^, we next investigated if this also applied to ET in thermoneutral conditions. The size of all analyzed fat depots was reduced similarly by ET in both temperatures (inguinal (i)WAT (−30%, Fig. 5A), epididymal (e)WAT (−40%, Fig. 5B), and BAT (−15%, Fig 5C). iWAT (Fig. 5A) and eWAT (Fig. 5B) mass were unchanged by housing temperature, while BAT amount was doubled in thermoneutrally-housed mice (Fig. 5C) supporting a recent study^20^. Thermoneutrality lowered basal glucose uptake in BAT by 85% with no effect observed in WAT depots (Suppl. Fig. 4a). ET did not alter basal glucose uptake in any of the analyzed adipose tissue depots (Suppl. Fig. 4a). ET in 22°C increased insulin-stimulated glucose uptake in iWAT, (+45%, Fig. 5D) and eWAT (+70%, Fig. 5E), but not in BAT (Fig. 5F). This response was unaffected by thermoneutral housing in eWAT but blunted by 30°C housing in iWAT. As seen for skeletal muscle, the differences in iWAT of ET-induced enhanced insulin action were ascribed to housing temperature rather than training volume ET also enhanced insulin action in iWAT in paired 22°C mice (Fig. 5D+E). 30°C housing led to an 80% reduction in insulin-stimulated glucose uptake in BAT compared to 22°C with no apparent effect of ET (Fig. 5F), and therefore the BAT from paired 22°C ET mice was not analyzed. To mechanistically explain the altered improvement in glucose uptake in adipose tissue, we analyzed the expression of glucose handling and insulin sensitive proteins. In iWAT, ET at 22°C led to increased HK II (+105%), GLUT4 (+105%), GS (+75%), and PDH (+95%), while only GLUT4 (+50%, p=0.094) and GS (+35%) increased after ET in 30°C (Fig. 5G). Like in muscle, HK II and PDH are thus potentially involved in the mechanisms behind the observed differences in insulin-stimulated glucose uptake in WAT. However, although showing tendencies for increased protein expression for most proteins investigated (incl. HK II and PDH), no significant effects were observed in the paired 22°C ET mice in iWAT (Suppl. Fig. 4b). Canonical insulin-stimulated signaling in iWAT was not affected by ET or housing temperature (data not shown), and thus could not explain the differences in insulin-stimulated glucose uptake. For BAT, only HK II (+45%) increased with ET in 22°C. In accordance with the lower insulin-stimulated glucose uptake in BAT at 30°C, we observed lower HK II (−90%), GLUT4 (−20%), and PDH (−30%) protein expression, while GS was unchanged (Fig. 5H).

**Figure 5:**
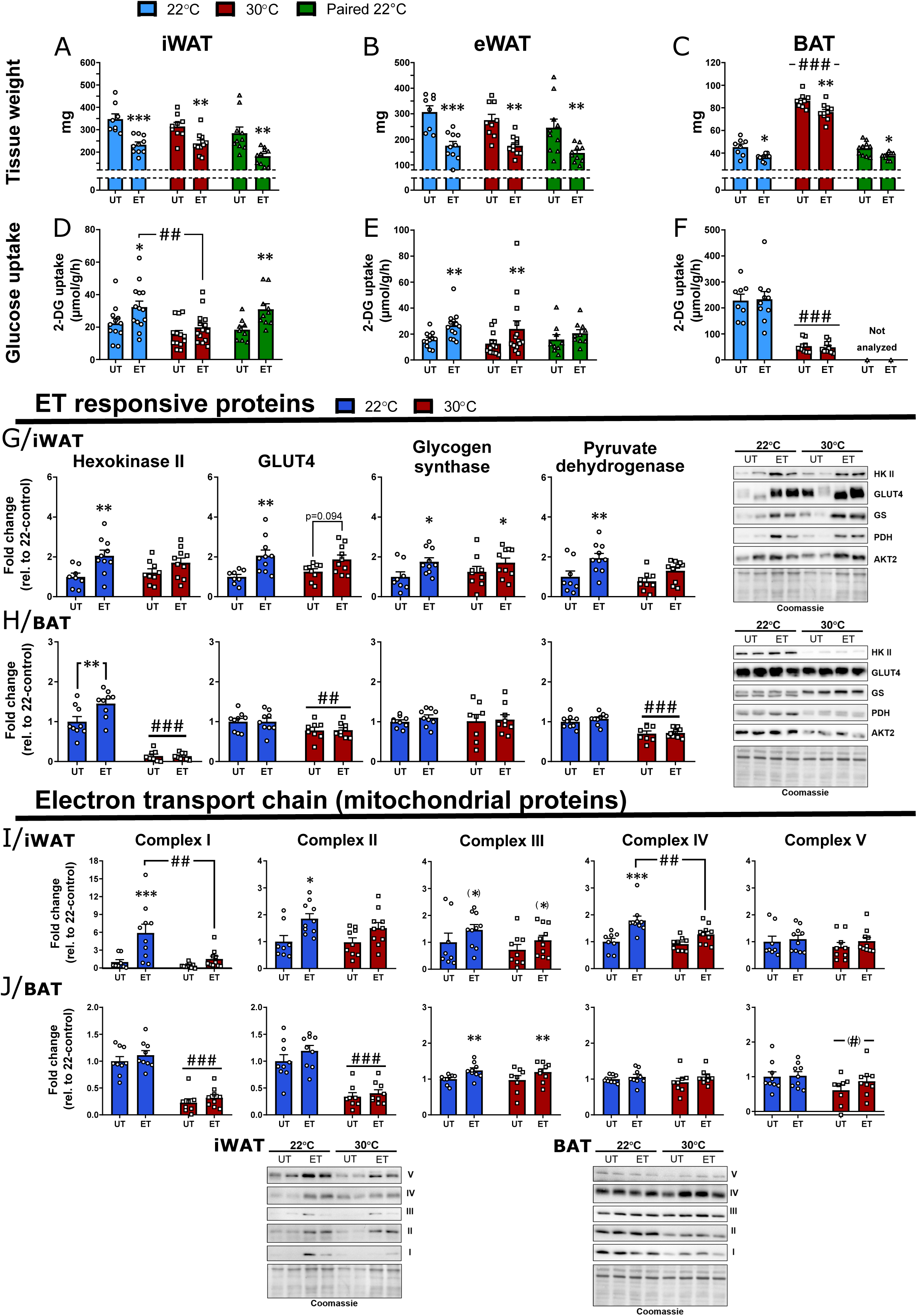
Exercise training improves insulin-stimulated glucose uptake in white adipose tissue to a greater extent when performed in ambient temperature. (A-C): Effect of voluntary wheel running exercise training (ET) on fat depot weight (iWAT (A), eWAT (B), and BAT(C)) at 22°C, 30°C, and paired 22°C, UT (untrained) and ET respectively. n=8-10. Effect of ET within temperature; * p<0.05, ** p<0.01, *** p<0.001. Main-effect of temperature; ### p<0.001. (D-F): Effect of ET on insulin-stimulated glucose uptake in 22°C, 30°C, and paired 22°C in iWAT (D), eWAT (E), and BAT (F). n=10-14. Effect of ET within temperature; * p<0.05, ** p<0.01. Effect of temperature as indicated with lines; ## p<0.01, ### p<0.001. (G-J): Effect of ET on training responsive proteins in iWAT (G) and BAT (H) at 22°C and 30°C, and the effect of ET on subunits of the electron transport chain of the mitochondrion in iWAT (I) and BAT (J) at 22°C and 30°C. Representative blots are shown as indicated. n=8-10. Effect of ET within temperature; * p<0.05, ** p<0.01, *** p<0.001. Main-effect of temperature within UT and ET groups; ## p<0.01, ### p<0.001. Parenthesis indicates p<0.1. Data are presented as mean ± SEM incl. individual values where applicable.

Because adipose tissue phenotypes are highly sensitive to temperature^34,35^ and ET has been reported to induce adipose tissue browning^36,38^, we analyzed gene expression of proteins involved in thermogenesis and mitochondrial uncoupling in iWAT and BAT. As expected, gene expression of proteins involved in thermogenesis (*Ucp1, Cidea, Prdm16, and PGC-1α*) were all downregulated by thermoneutral housing in all depots investigated (Suppl. Fig. 4c). These genes were largely unaffected by ET in both temperatures, likely due to the fact that the running wheels were locked for 24 hours prior to the terminal experiment. Indeed, ET increased protein content of four of five complexes of the electron transport chain in iWAT at 22°C in ET mice, but not at 30°C (Fig. 5I). Thus, paired 22°C ET only led to a significant increase in complex 3. Thus, both reduced running distance at thermoneutrality as well as temperature seem to underlie the differences in molecular adaptation to ET in iWAT. In BAT, only complex III increased (+25%) with voluntary ET and this occurred at both housing temperatures (Fig. 5J). Thermoneutral housing reduced complex I (−75%) and II (−65%) in BAT compared with 22°C (Fig. 5J). With strikingly no or only little effect observed of ET in BAT at either temperature and therefore likely not a key target for the observed phenotype in 30°C-housed mice, paired 22°C ET BAT was not investigated.

Despite having diminished molecular training adaptations in iWAT of the paired 22°C ET mice, increased insulin-stimulated glucose uptake was still observed in both ET-groups in iWAT of 22°C housing. Therefore, other unexplored or unknown mechanisms must underlie the observed effect on glucose metabolism in iWAT after ET. It emphasizes the importance of housing temperature when performing exercise training studies investigating fat depots and metabolism.

### Thermoneutral housing supersedes the effect of exercise on gut microbiome composition

Metabolic health has during recent years been shown to be under strong influence by the gut microbiome (GM)^39^, and we therefore tested if housing temperature affects ET-induced gut microbiome adaptations.

GM diversity following the different interventions is visualized in a PCA plot (Fig. 6A). Both UT and ET mice showed significant differences in GM composition with distinct separation depending on housing temperature (R = 0.21, p=0.001). Forty-six phylotypes with a cumulative relative abundance of up to 14%, differed significantly between the four experimental groups (Fig. 6B and C). These primarily belonged to family *Muribaculaceae* (40 phylotypes), a dominant bacterial group in the mouse gut^40^, followed by family *Lachnospiraceae* (3 phylotypes), species *Bacteroides uniformis* (2 phylotypes) and genus *Mobilisprobacter* (1 phylotype) (Fig. 6B and C).

**Figure 6:**
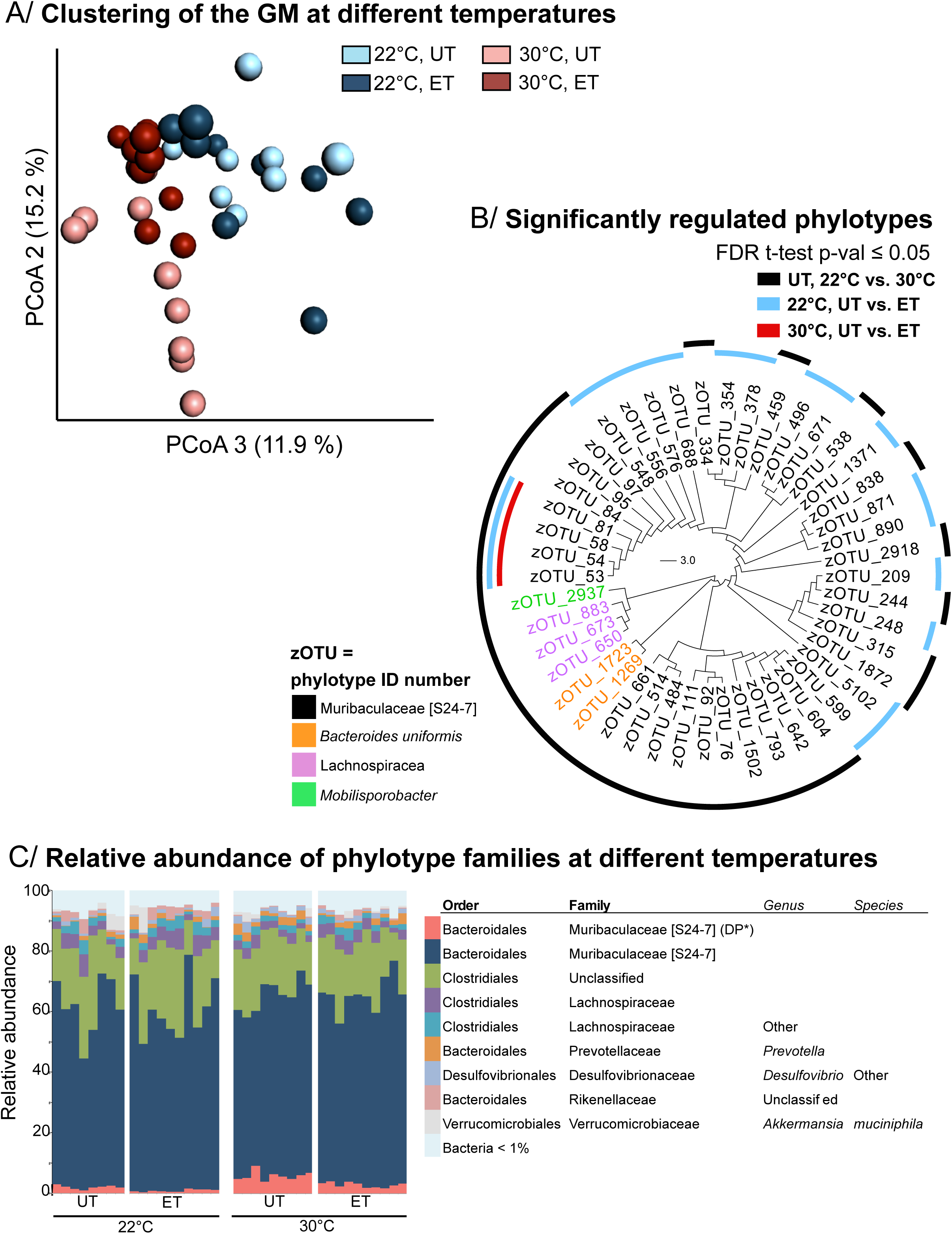
Thermoneutral housing supersedes the effect of exercise on gut microbiome composition. (A) Housing mice at 30°C markedly alters gut microbiome composition with minor effect of exercise training. Bray-Curtis distance based PCoA plot of cecal 16S rRNA gene (V3-region) amplicons (zOTU level). The groups are separated by the colors indicated in the figure. Adonis test determined differences between experimental groups [R^2^ = 0.21, *p*-val = 0.001], where a clear separation between mice housed at 22°C and 30°C was observed. n= 8-10. (B) Specific phylotypes (summarized with zOTUs) were differentially regulated by housing temperature and exercise training. *t*-test (*FDR p*-val ≤ 0.05). n= 8-10. (C) Relative abundance of zOTUs summarized to the species level as indicated. n= 8-10.

With only four phylotypes changed with ET at 30°C and 19 changed in 22°C, the effect of housing temperature superseded the effect on GM compared to ET. However, four phylotypes, members of *Muribaculaceae*, were affected by both ET as well as housing temperature (Fig. 6B). These phylotypes have not been described at species level yet and additional studies are needed to define the role of these bacteria in relation to metabolism.

Significant positive and negative correlations between 676 phylotypes (a cumulative abundance of 10.2% of the entire dataset) and insulin-stimulated glucose uptake of all analyzed tissues (apart from the heart) and fasting blood glucose were observed (Suppl. Fig.5). *Bacteroidaceae* (*B. uniformis*), *Porphyromonadaceae* (*P. distonis*), and *Ruminococcaceae* (*Oscillospira* spp.) and most *Bifidobacteriaceae* and *Coreobacteriaceae* members were all negatively correlated to these parameters (Suppl. Fig. 5).

The above data demonstrate that although ET alters the abundance of specific phylotypes at different housing temperatures, housing temperature of 30°C *per se* also causes a remarkably modification of the GM composition.

## Discussion

The major finding in the current study was that housing temperature significantly alters systemic metabolic as well as molecular adaptations to voluntary wheel running exercise training in mice. In recent years, it has become evident that housing temperature markedly affects mouse metabolism and this complicates the translatability to humans. Notably, at ambient housing temperature over one-third of total energy expenditure in mice is cold-induced thermogenesis^41^. In contrast, cold-induced thermogenesis contributes a very small fraction to total energy expenditure in humans^42^. Increasing housing temperatures (27°C - 30°C) improves the metabolic similarity between humans and mice and has been suggested to be a better housing strategy^9,10,26,27,33^. To the best of our knowledge, this is the first study to show that housing temperature markedly influences exercise training adaptations, clearly demonstrating that housing temperature is an important consideration when investigating such parameters in mice.

Most remarkably, the ET response on glucose metabolism was reduced by thermoneutral housing. ET at ambient temperature led to an increased glucose tolerance and improved insulin-stimulated glucose uptake in skeletal muscle in agreement with many previous reports^43–51^. However, this effect was absent in thermoneutrally housed mice. With regards to glucose tolerance, this could be ascribed to reduced running volume in thermoneutrally housed mice because paired 22°C mice, that ran the same distance as the 30°C mice, also did not improve glucose tolerance with training. However, the paired 22°C mice still improved insulin sensitivity as suggested by a reduced glucose-stimulated insulin response during the glucose tolerance test as well as improved insulin-stimulated skeletal muscle and iWAT glucose uptake. In contrast to glucose tolerance, the blunted ET-induced enhanced insulin-stimulated glucose uptake observed in 30°C housed mice, could be solely ascribed to housing temperature and not training volume. In addition, while 22°C-housing led to mild cold stress during day-time, the observed changes (running distance e.g.) were not due to overheating of 30°C-housed mice. Our finding that the voluntary wheel running model is less efficient in improving metabolic status in mice housed under thermoneutral conditions, could indicate that this is in fact not a good choice of housing condition when investigating molecular events underlying the metabolic benefits of exercise. On the other hand, at 22°C, the mouse has an extraordinarily high running volume, hardly mimicking a human exercise intervention. Combined with mild cold stress, such excessive exercise regimes could mask or intensify potential effects when investigating a genetic model or pharmacological compound. This has indeed been observed for many other molecular mechanisms, where conclusions drawn from mice housed at 22°C have been completely changed when investigated at thermoneutrality^17,19,21–24,52,53^.

Another major finding of our study was that the molecular adaptations of key exercise responsive proteins were markedly altered by thermoneutrality, and this was not due to lower running volume. In triceps muscle, hexokinase II (responsible for upholding the glucose gradient across the membrane by phosphorylating entering glucose) and pyruvate dehydrogenase (converts pyruvate to acetyl-CoA connecting the glycolysis and the Krebs cycle) were only significantly upregulated by ET in ambient temperature, not thermoneutrality. This was also apparent for subunits of the electron transport chain in the mitochondria. Interestingly, these differences in mitochondrial adaptations in muscle and fat depots were not reflected in differences in improvements in exercise performance, as running capacity increased equally in all ET groups.

Metabolic ET-induced improvements are often associated with reduced adiposity^54,55^. In our study, ET reduced body-fat at both housing temperatures, although this was observed to a lesser extent at thermoneutrality. Interestingly, our paired running group elucidated that this difference was a cause of housing temperature and not training volume. A better effect of ET on reducing adiposity at 22°C is likely due to the much higher metabolic demand on mice at 22°C that, because of mild cold stress, exhibit increased energy expenditure as has been described previously in UT mice^56^. Our metabolic measurements of mice during voluntary wheel running at different temperatures showed, that mice running at ambient temperature have much higher oxygen consumption, and thereby also increased energy usage. This could be a contributing factor for a lower body fat percentage seen in trained mice at 22°C compared to 30°C. Virtue and colleagues (2012) have found somewhat opposite results showing that mice running at 28°C displayed higher energy usage per wheel turn compared to 21°C^57^. However, in that study, the mice were not habituated to thermoneutral temperatures prior to the experiment, which might explain the discrepancy.

In addition to a reduced metabolic rate, mice housed at 30°C did not increase their RER during the dark cycle. A lower RER is indicative of higher relative contribution of fat oxidation. Taken together with the observed higher FFA and TG levels of 30°C-housed mice (in agreement with a previous report^20^), the data from current study shows that fat metabolism might also be highly affected by housing temperatures and warrants further investigations.

Although not a key objective of the study, we found generic differences between housing temperatures in our untrained mice that to the best of our knowledge have not previously been documented. An important finding in our study was that the mouse gut microbiome was remarkably affected by housing temperature with minimal effect of ET. The small effect size of ET on GM contrasts previous observations^58,59^. The effect of temperature is important as the gut microbiome has been shown to affect several functions in physiology, e.g. glucose metabolism^60–62^, and has recently been associated with muscle function^63^. This effect of housing temperature alone has not to our knowledge been clearly demonstrated previously.

Increasing housing temperature also led to a ~10% reduction in heart mass, suggesting that ambient housing leads to cardiac hypertrophy, likely due to a higher cardiac stress as indicated by twice as high heart rate at 22°C (600bpm) compared to 30°C (300bpm) housed mice^15,64,65^.

In addition to a lower fasting blood glucose of thermoneutrally housed mice as observed previously^19^, we also observed that thermoneutral housing lead to elevated insulin secretion following a glucose challenge. To what extent housing temperature alters pancreatic morphology or function is unknown and needs further investigation. Based on our findings, housing temperature may affect the outcome of studies investigating all of these processes, the interpretation of the results, and ultimately the translation to humans.

Contemporary biomedical research is using proteomic analysis to comprehensively explore the global regulations to map the beneficial changes that occurs with exercise training^49,66–68^. Such studies will need to be followed up by hypothesis-driven research genetically manipulating or pharmacologically inhibiting/activating a pathway of interest in order to elucidate the mechanistic role for a given exercise training-regulated protein or process. Considering the optimal housing condition for such studies might increase the translatability and clinical relevance for humans.

## Conclusion

In conclusion, we show that numerous training adaptions are influenced by housing temperature; the majority of which was not ascribed to a lower voluntary running volume in thermoneutrally-housed mice. Our findings highlight that organismal and molecular adaptations to exercise training in mice depend upon housing temperature and that housing temperature is important to consider when using mice as an experimental model.

## Additional information

### Competing interests

None declared

### Author contributions

S.H.R., L.S., and E.A.R. conceptualized and designed the study. S.H.R. and L.S. conducted the experiments, performed the laboratory analysis, analyzed the data, and wrote the manuscript. C.H.O., I.K., M.A., L.L.V.M., W.K., J.L.C.M., D.S.N., and Z.G.H. all took part in conducting the experiments, performing laboratory analysis and/or interpreting the data. All authors commented on and approved the final version of the manuscript. L.S. is the guarantor of this work and, as such, has full access to all the data in the study and takes responsibility for the integrity of the data and the accuracy of the data analyses.

### Funding

L.S. and E.A.R. were supported by the Danish Council for Independent Research, Medical Sciences (grant DFF-4004-00233 to LS, grant 6108-00203 to EAR); The Novo Nordisk Foundation (grant 10429 to EAR, grant NNF16OC0023418 and NNF18OC0032082 to LS). L.L.V.M. was supported by the PhD fellowship from The Lundbeck Foundation (grant 2015-3388 to LLVM).

## Acknowledgements

We acknowledge the skilled technical assistance of Betina Bolmgren and Irene Bech Nielsen (Molecular Physiology Group, Department of Nutrition, Exercise and Sports, University of Copenhagen, Denmark).

## Material and methods

### Animals

10-week old female C57BL/6J mice (Taconic, Lille Skensved, Denmark) were maintained on a 12:12-h light-dark cycle and received standard rodent chow diet (Altromin no. 1324; Chr. Pedersen, Denmark) and water ad libitum with nesting materials. All experiments were approved by the Danish Animal Experimental Inspectorate (Licence; 2016-15-0201-01043). Mice were randomly assigned to ambient temperature (22°C±1°C) or thermoneutrality (30°C±1°C) in different rooms in the same animal facility. After a 7-10 day acclimatization period, mice were pair-housed and housed with or without free access to running wheels for 6 weeks. Running distance was recorded for each cage twice weekly. Core temperature was measured with a rectal thermometer at 1:00pm (light period) and 9:00pm (most active dark period). For paired exercise training mice were housed at ambient temperature with wheels that were locked from 00.00am-5:00pm. Running wheels were locked 24 hrs before glucose tolerance tests and terminal procedures to avoid any residual effects of acute exercise.

### Maximal running capacity

Mice were acclimated to the treadmill three times (10 min at 0.16 m/s) within a week prior to the maximal running tests. The maximal running test started at 0.16 m/s for 300 s with 15° incline, followed by a continuous increase (0.2 m/s) in running speed every 60s until exhaustion (Treadmill TSE Systems, Germany).

### Body composition

Total, fat and lean body mass were measured weekly by nuclear magnetic resonance using an EchoMRI™(USA).

### Metabolic chambers

After a 3-day acclimation period in the metabolic cages, oxygen consumption, ambulant activity (beam breaks), food intake and running distance/speed were measured by indirect calorimetry in a CaloSys apparatus during at least 2 days (TSE Systems, Bad Homburg, Germany). To test the acute effects of thermoneutrality, a group of mice were housed in the metabolic chambers at 22°C for 3 days followed by an increase in the temperature to 30°C for 3 days. All mice were single-housed during housing in metabolic cages.

### Glucose tolerance test

Glucose (2.0 g/kg) was intraperitoneally injected into 5-hr-fasted (fasting from 7:00AM) mice. Blood was collected from the tail vein at time points 0, 20, 40, 60, 90 and 120 min and analyzed for glucose using a glucometer (Bayer Contour; Bayer, Münchenbuchsee, Switzerland). At time point 0 and 20 min, insulin was analyzed in duplicates in plasma (#80-INSTRU-E10; ALPCO Diagnostics).

### *In vivo* insulin-stimulated ^3^H-2-DG uptake

To determine 2-deoxyglucose (2-DG) uptake in muscle, [^3^H]2-DG (Perkin Elmer) was injected retro-orbitally in a bolus of saline containing 66.7 μCi/mL [^3^H]2DG corresponding to ~9-10 μCi/mouse (6 μL/g body weight) in chow. The injectate also contained 0.3 U/kg body weight insulin (Actrapid; Novo Nordisk, Bagsværd, Denmark) or a comparable volume of saline. Prior to stimulation, mice were fasted for 3 h from 07:00am and anaesthetized (intraperitoneal injection of 7.5 mg pentobarbital sodium/100 g body weight) for 15 min. Blood samples were collected from the tail vein immediately prior to insulin or saline injection and after 5 and 10 min and analyzed for glucose concentration using a glucometer (Bayer Contour; Bayer, Münchenbuchsee, Switzerland). After 10 min, all tissues were excised, weighed (fat depots only), and quickly frozen in liquid nitrogen and stored at −80°C until processing. Blood was collected by punctuation of the heart, centrifuged and plasma frozen at −80°C. Plasma samples were analyzed for insulin concentration and specific [^3^H]2DG tracer activity. Tissue specific 2DG uptake was analyzed as previously described (Fueger et al. 2004; Raun et al. 2018).

#### Plasma analysis

Plasma insulin concentration was analyzed in duplicates in plasma (#80-INSTRU-E10; ALPCO Diagnostics). Plasma triacyglyceride (TG) was analyzed in duplicates in plasma (#Triglycerides CP, Horiba ABX). Plasma free fatty acids (FFA) were analyzed in duplicates in plasma (#NEFA C ACS-ACOD, Wako Chemicals).

#### Tissue processing

Muscles were pulverized in liquid nitrogen and homogenized 2 × 0.5 min at 30 Hz using a TissueLyser II bead mill (Qiagen, USA) in ice-cold homogenization buffer (10% glycerol, 1% NP-40, 20 mM sodium pyrophosphate, 150 mM NaCl, 50 mM HEPES (pH 7.5), 20 mM β-glycerophosphate, 10 mM NaF, 2 mM phenylmethylsulfonyl fluoride (PMSF), 1 mM EDTA (pH 8.0), 1 mM EGTA (pH 8.0), 2 mM Na3VO4, 10 μg mL^−1^ leupeptin, 10 μg mL^−1^ aprotinin, 3 mM benzamidine). Following end-over-end rotation for 30 min at 4°C, the samples were centrifuged (10,000rpm) for 20 min at 4°C. Thereafter, the supernatants (clear lysate) were collected and stored at −80°C. The latter two steps (centrifugation and lysate collection) were performed three times in adipose tissue to avoid contamination of fatty acids.

#### Immunoblotting

Lysate protein concentrations were measured using the bicinchoninic acid (BCA) method with bovine serum albumin (BSA) as standard. Total protein and phosphorylation levels of relevant proteins were determined by standard immunoblotting techniques loading equal amounts of protein. The primary antibodies used are presented in Table 1. Polyvinylidene difluoride membranes (Immobilon Transfer Membrane; Millipore) were blocked in Tris-buffered saline (TBS)-Tween 20 containing 2% milk protein for 5 min at room temperature. Membranes were incubated with primary antibodies overnight at 4°C, followed by incubation with horseradish peroxidase-conjugated secondary antibody for 45 min at room temperature. Coomassie brilliant blue staining was used as a loading control ^69^. Bands were visualized using the Bio-Rad ChemiDoc MP Imaging System and enhanced chemiluminescence (ECL+; Amersham Biosciences).

**Antibody table 1.**
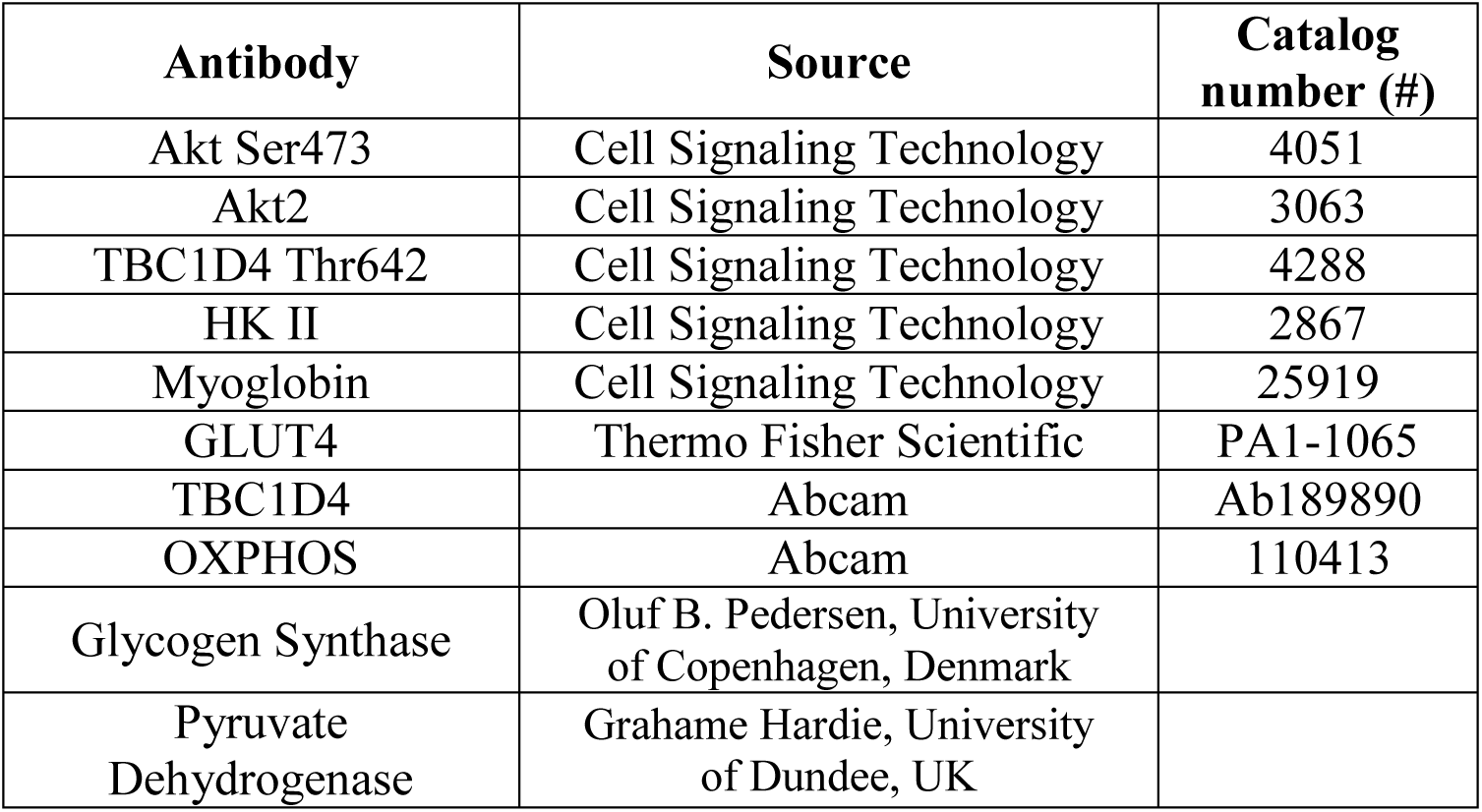

#### qPCR analysis

Total RNA was extracted from BAT, iWAT and eWAT depots using TRI reagent (T9424, Sigma-Aldrich) followed by isolation using RNeasy Mini Kit (74106, Qiagen). Reverse transcription was carried out on 1000 ng RNA using the High Capacity cDNA Reverse Transcription kit (4368814, Applied Biosystems). Gene expression was determined based on real-time quantitative PCR using SYBR green (PP00259, Primerdesign). The data was analyzed with the ΔΔCT method and normalized to the housekeeping gene 36b4. All primers are listed in Table 2.

**Primer table 2.**
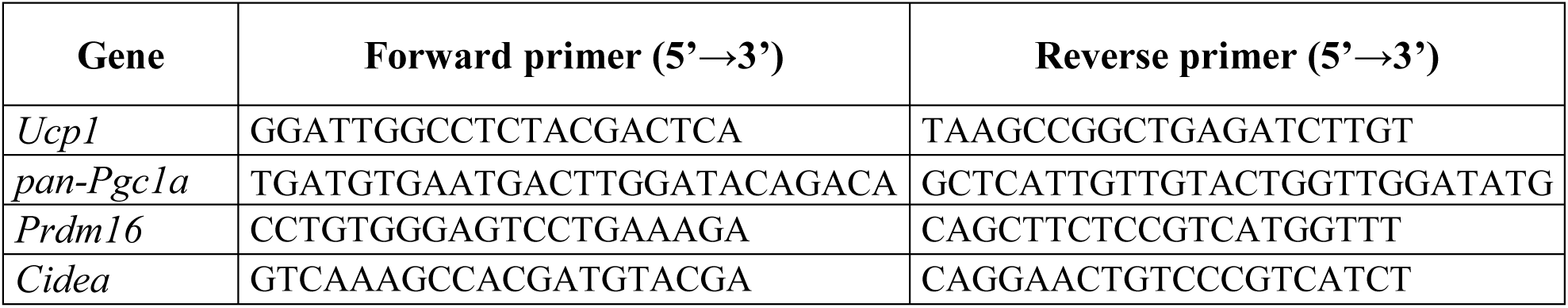

### Microbiota analysis

#### Samples collection, processing and DNA extraction

Caecum fecal samples were collected from sedated mice during the terminal experiment with R.O. injections. Approximately 200 mg of the caecal content were used for DNA extraction using the PowerSoil® DNA Isolation Kit (MOBIO Laboratories, Carlsbad, CA, USA), following the instructions of the manufacturer, but with minor modifications. Briefly, prior DNA extraction, samples were placed into the PowerBead tubes and heat treated at 65°C for 10 min and then at 95°C for 10 min. Subsequently, solution C1 was added and bead-beating performed in FastPrep (MP Biomedicals, Santa Ana, CA, USA) using 3 cycles of 15 s each, at a speed of 6.5 m s^−1^. The remaining DNA extraction procedure followed the manufacturer’s instructions.

#### High-throughput 16S rRNA gene amplicon sequencing

Gut microbiome composition was determined by high-throughput 16S rRNA gene amplicon sequencing. The primers designed with adapters Nextera Index Kit® (Illumina, CA, USA) targeted the V3 region (~190 bp) and the library preparation, purification and sequencing were performed as previously described ^70^. Briefly, the amplification profile (1^st^ PCR) followed: Denaturation at 95°C for 2 min; 33 cycles of 95°C for 15 s, 55°C for 15s and 68°C for 40 s; followed by final elongation at 68°C for 5 min, while barcoding (2^nd^ PCR) was performed at 98°C for 1 min; 12 cycles of 98°C for 10 s, 55°C for 20 s and 72°C for 20 s; elongation at 72°C for 5 min. The amplified fragments with adapters and tags were purified and normalized using custom made beads, pooled and subjected to 150 bp pair-ended NextSeq (Illumina, CA, USA) sequencing.

Sequencing of the 16S rRNA gene (V3-region) amplicons yielded 3,434,893 high quality reads (mean sequence length of 183 bp) and the number of reads per sequenced sample varied from 50,140 to 141,905 with an average of 85,872 (SD 20,246).

#### Processing of high throughput sequencing data

The raw dataset containing pair-ended reads with corresponding quality scores were merged and trimmed using the following settings, -fastq_minovlen 100, -fastq_maxee 2.0, -fastq_truncal 4, -fastq_minlen 130. De-replicating, purging from chimeric reads and constructing *de-novo* zero-radius Operational Taxonomic Units (zOTU) was conducted using the UNOISE pipeline ^71^ coupled to the EZtaxon 16S rRNA gene collection as a reference database ^72^.

### Statistical Analyses

The data are expressed as mean ± SEM and individual data points (when applicable) and analyzed using GraphPad Prism 8. Statistical tests were performed using paired/non-paired t-tests or repeated/no-repeated two-way ANOVA as applicable. Multiple repeated Two-way ANOVAs were performed in analyses including all experimental groups testing for the effect of temperature within training groups or the effect of exercise training within each temperature. Sidak post-hoc test was performed when ANOVA revealed significant main effects and interactions. Microbiome analyses; For downstream analyses the dataset (based on zOTUs phylotypes) was subsampled with 50,000 reads. Principal Coordinates Analysis (PCoA) based on Bray-Curtis distances (based on 10 distance metrics and determined by 10 subsampled zOTU tables) and differences between experimental groups were evaluated using analysis of variance on distance matrices (Adonis). Differences in relative distribution among phylotypes were determined through Student’s *t*-test, while correlations of phylotypes with measurements for insulin-stimulated glucose uptake were determined with Pearson Correlation Coefficients. These analyses were bootstrapped with 100 permutations and *p*-values corrected for Type I error with False Discovery Rate (FDR).

## Figure legends (Appendix, supplementary data)

**A1: For figure 1 and 2.**
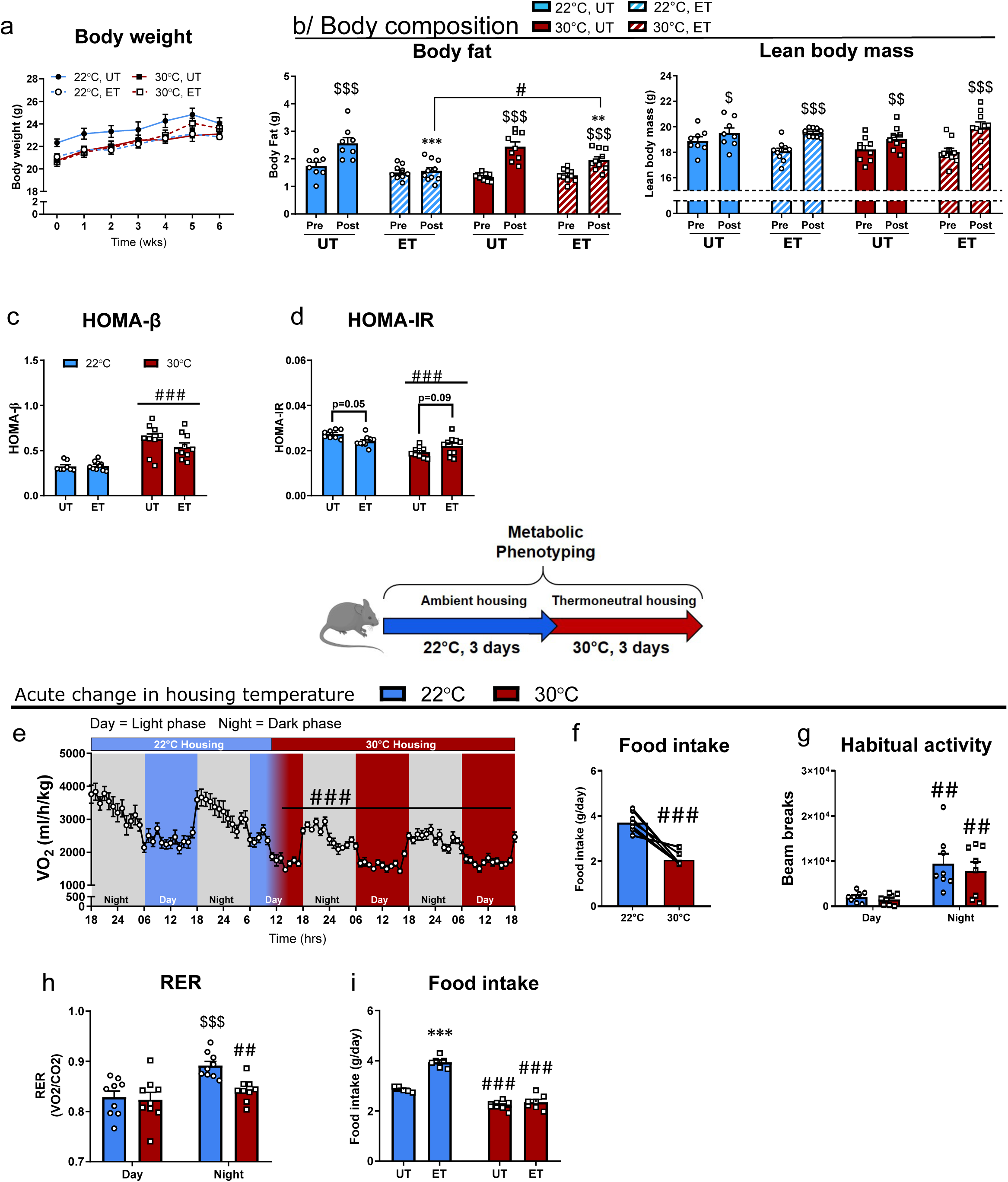
(a) The effect of housing temperature and ET in 22°C and 30°C on bodyweight (gram). n=8-10. (b) The effect of housing temperature and ET in 22°C and 30°C on body fat (gram) and lean body mass (gram). n=8-10. Effect of time within group; $ p<0.05, $$$ p<0.001. Effect of temperature within ET-groups (post); # p<0.05. Effect of ET within temperature; * p<0.05, *** p<0.001. (c-d) HOMA-β (beta-cell sensitivity) and HOMA-IR (whole body) measured from basal plasma glucose and insulin. n=8-10. Effect of temperature; ### p<0.001. (e-h): The effect of acute change of temperature from 22°C to 30°C on oxygen uptake (VO2) food intake, ambulant activity (2 consecutive days), and RER. n=9. Effect of time; ## p<0.01, ### p<0.001. (i): Food intake in ET mice housed in metabolic chambers. Effect of ET; *** p<0.001. Effect of temperature; ### p<0.001 (n=5-8). Data are presented as mean ± SEM incl. individual values where applicable.

**A2: for Figure 3:**
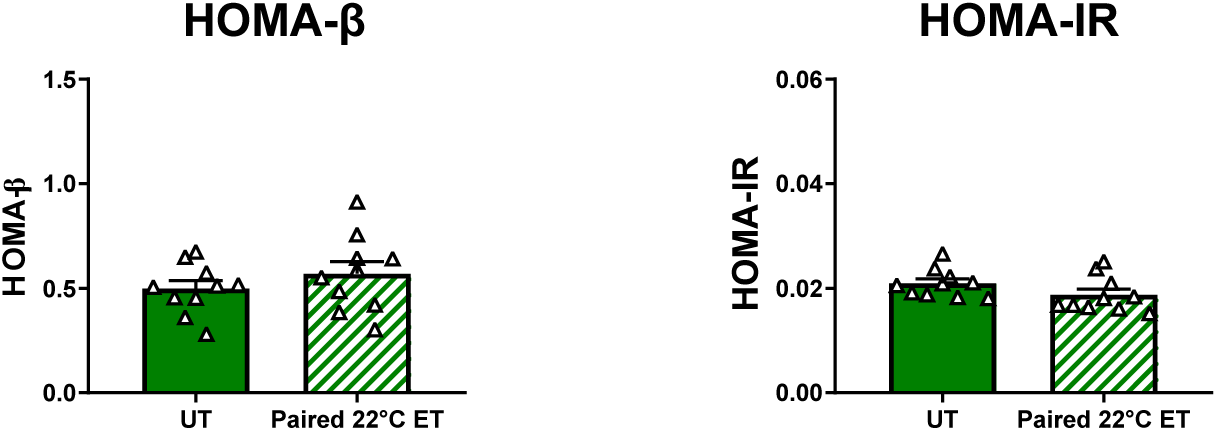
HOMA-β (beta-cell sensitivity) and HOMA-IR (whole body) measured from basal plasma glucose and insulin. n=10 Data are presented as mean ± SEM incl. individual values where applicable.

**A3: for Figure 4.**
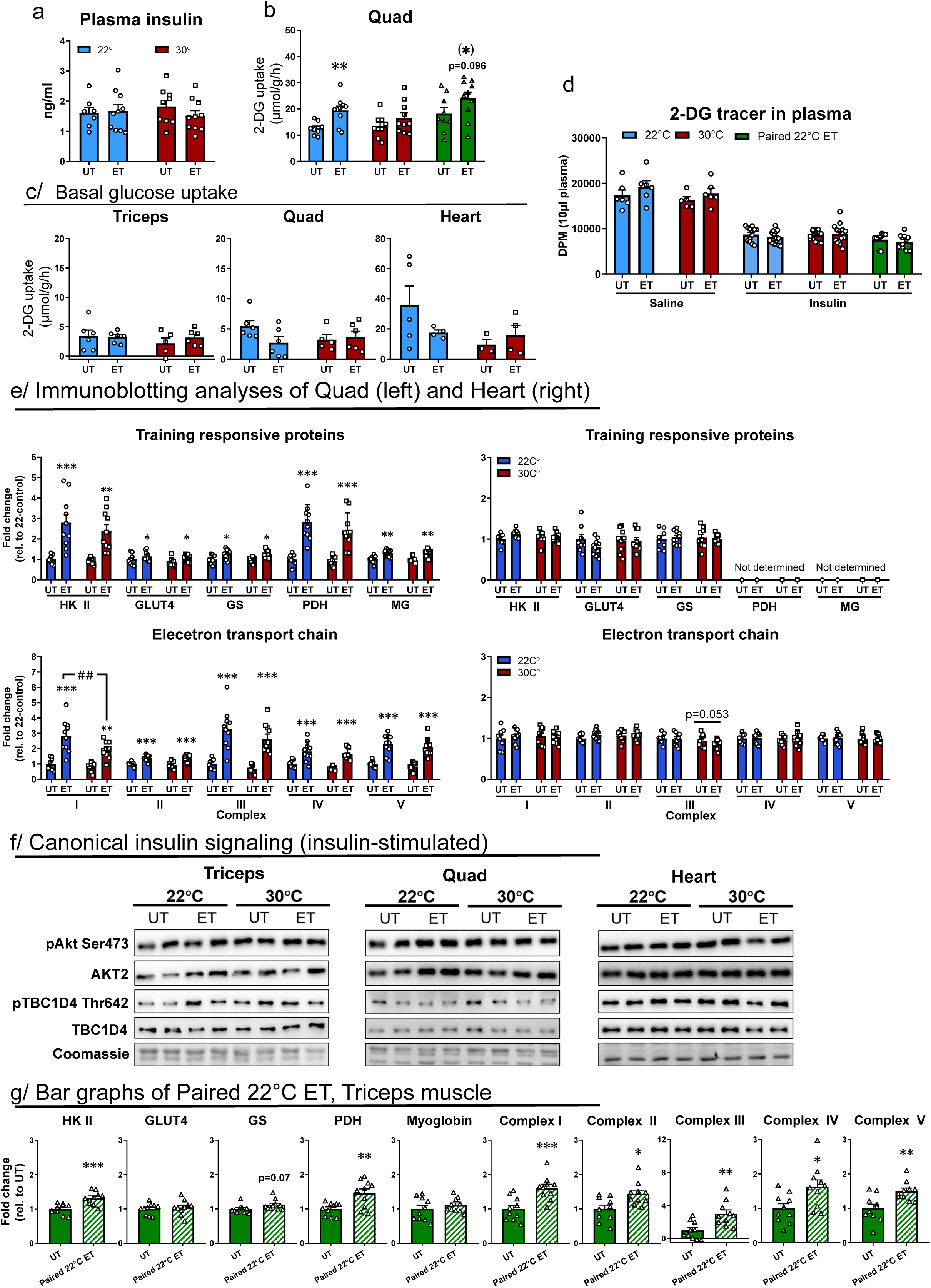
(a) Plasma insulin after retro-orbital insulin injection (at min 10). n=8-10, Two-way ANOVA. (b) Effect of ET on insulin-stimulated glucose uptake in 22°C, 30°C, and paired 22°C in skeletal muscle (m. quadriceps). n=8-10. Effect of ET within temperature; * p<0.05. (c) Basal glucose uptake in all experimental groups. n=6 (d) 2-DG plasma tracer counts of all experimental groups (e) Effect of ET on training responsive proteins and subunits of the electron transport chain of the mitochondrion in triceps muscle at 22°C and 30°C. n=8-10. Effect of ET within temperature; * p<0.05, ** p<0.01, *** p<0.001. Effect of temperature within UT or ET groups; ## p<0.01. (f) Representative blots of canonical insulin signaling in all muscles investigated. (g) Bar plots of paired 22°C ET immunoblotting analyses. Representative blots can be seen in figure 4. n=10. Effect of ET; * p<0.05, ** p<0.01, *** p<0.001. Data are presented as mean ± SEM incl. individual values where applicable.

**A4: for Figure 5.**
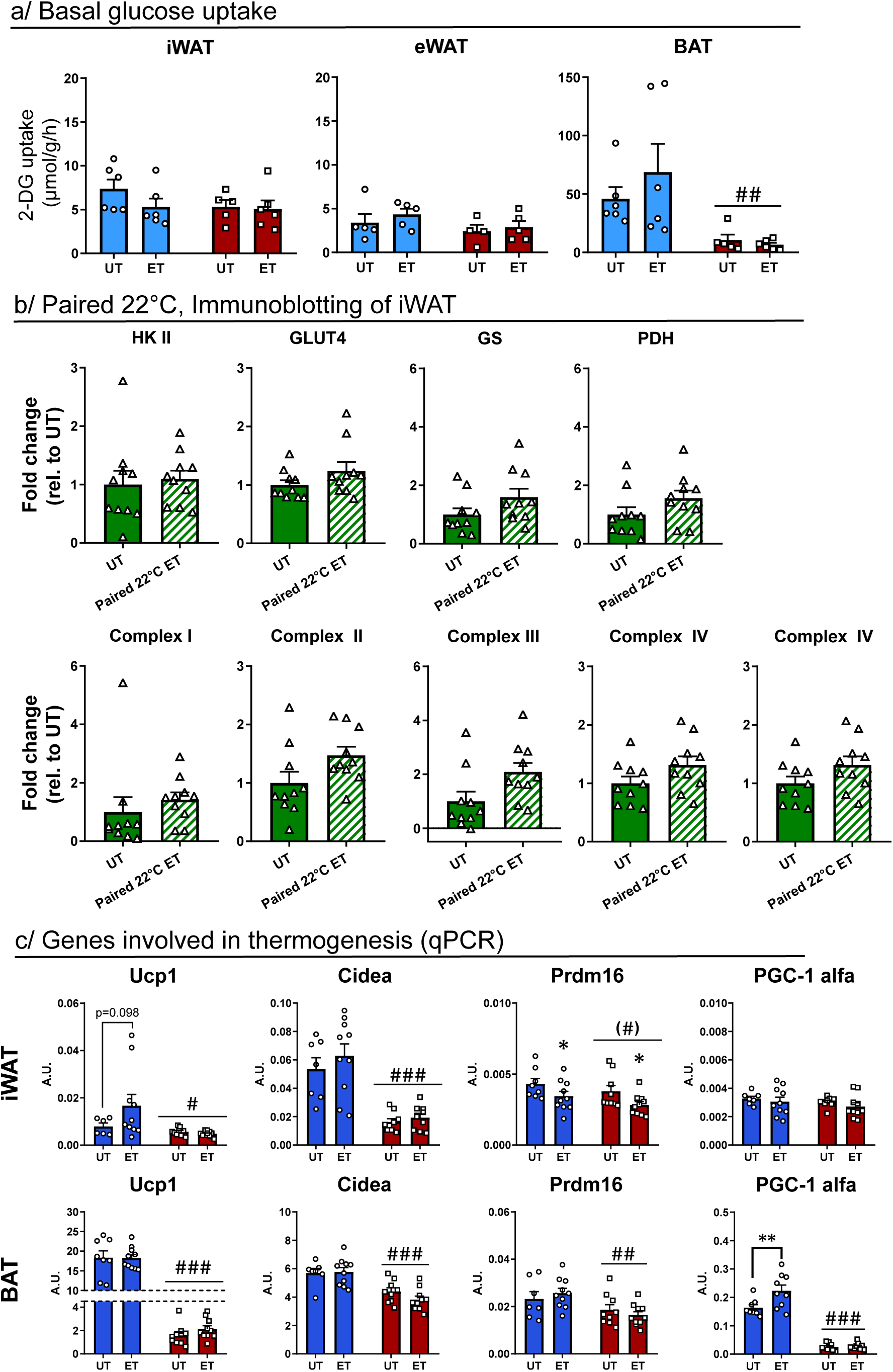
(a) Basal glucose uptake in all experimental groups. n=6 (c) Bar plots of immunoblotting analyses of paired 22°C ET of iWAT. (b) Thermo-regulatory genes in iWAT and BAT depots. n=7-10. Effect of ET within temperature; * p<0.05. Effect of temperature within UT or ET groups; ## p<0.01, ### p<0.001, parenthesis; p<0.1. Data are presented as mean ± SEM incl. individual values where applicable.

**A5: for Figure 6:**
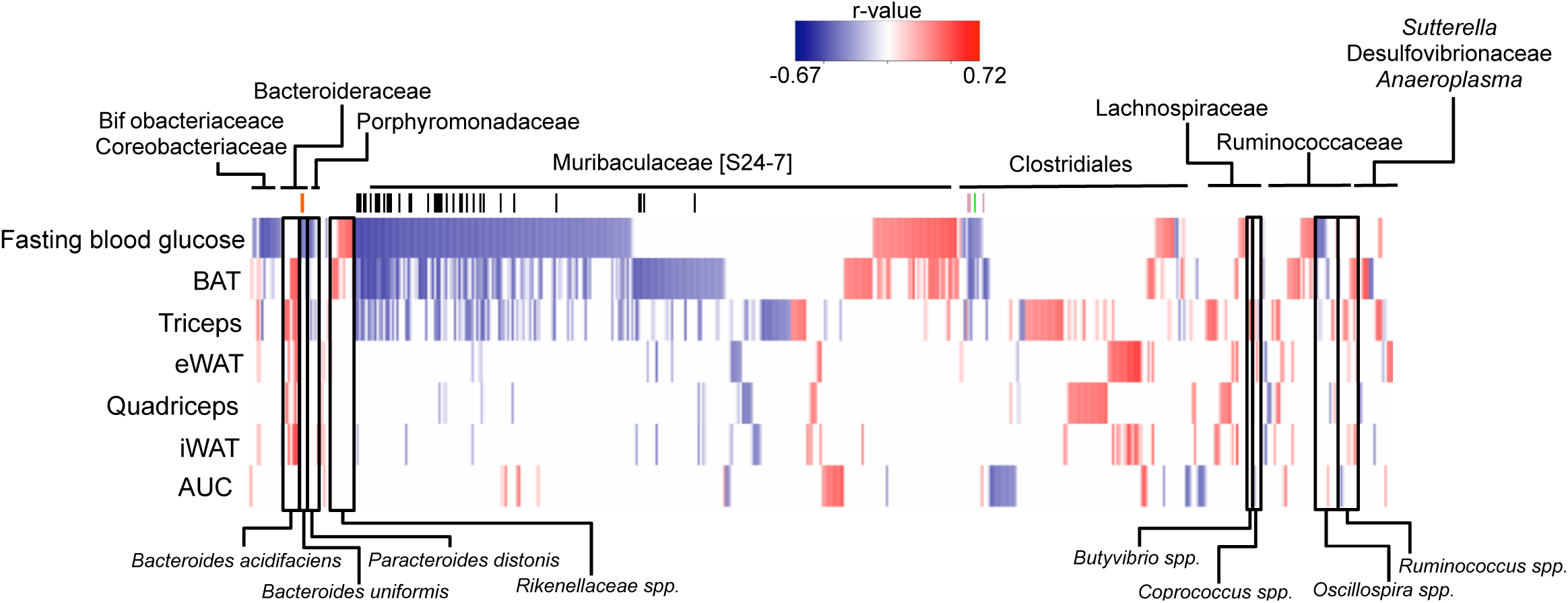
(A) Heat-map displaying Pearson Correlation Coefficients (FDR *p*-val ≤ 0.05) between measurements of insulin-stimulated glucose uptakes and the relative abundance of 676 GM phylotypes. Blue color indicates negative correlation. Red color indicates positive correlation.

